# FEVER: An interactive web-based resource for evolutionary transcriptomics across fishes

**DOI:** 10.1101/2024.03.22.586044

**Authors:** Jérôme Montfort, Francisca Hervas-Sotomayor, Aurélie Le Cam, Florent Murat

## Abstract

Teleost fish represent one of the largest and most diverse clades of vertebrates, which makes them great models in various research areas such as ecology and evolution. Recent sequencing endeavors provided high-quality genomes for species covering the main fish evolutionary lineages, opening up large-scale comparative genomics studies. However, transcriptomic data across fish species and organs are heterogenous and have not been integrated with newly sequenced genomes making gene expression quantification and comparative analyses particularly challenging. Thus, resources integrating genomic and transcriptomic data across fish species and organs are still lacking. Here, we present FEVER, a web-based resource allowing evolutionary transcriptomics across species and tissues. First, based on query genes FEVER reconstructs gene trees providing orthologous and paralogous relationships as well as their evolutionary dynamics across 13 species covering the major fish lineages, and 4 model species as evolutionary outgroups. Second, it provides unbiased gene expression across 11 tissues using up-to-date fish genomes. Finally, genomic and transcriptomic data are combined together allowing the exploration of gene expression evolution following speciation and duplication events. FEVER is freely accessible at https://fever.sk8.inrae.fr/.

## INTRODUCTION

Gene duplication and subsequent expression changes (e.g., sub/neo-functionalization) are thought to underlie many phenotypic differences across animal species (1–6). Teleost fish represent a unique opportunity to study this phenomenon as they nearly represent half of all vertebrate species and exhibit an outstanding phenotypic diversity due to their adaptation to a wide range of ecological niches. Notably, their common ancestor underwent a Whole Genome Duplication event (WGD) approximately 300 million years ago (called Ts3R) that provided new raw genetic material for natural selection to act on (7). Since then, intense genomic reshuffling occurred which probably facilitated the diversification of species and organs (8). Specific fish lineages, such as salmonids (9) and carps (10), underwent supplementary WGDs (called Ss4R and Cs4R, respectively). Additionally, fish species such as zebrafish and medaka represent major models in various research fields such as developmental biology, biomedical research, ecology and evolution. Recent sequencing endeavors provided high-quality genomes for species covering the main fish evolutionary lineages (8, 11). However, transcriptomic data across fish species and organs are heterogenous and have not been integrated with newly sequenced genomes making gene expression quantification and comparative analyses particularly challenging. Thus, tools allowing the exploration of gene evolutionary history and their expression profiles across species and organs in fish are still lacking. To fill this gap, we developed the FEVER web service which provides a user-friendly interface to rapidly explore evolutionary dynamics of genes of interest across fish species and tissues at the structural (gene gain and loss) and expression levels. To do so, FEVER reconstructs gene trees using methods inspired by landmark comparative resources (12–14) that provide orthologous and paralogous relationships as well as gene evolutionary dynamics across 13 species covering the major fish lineages (bowfin, spotted gar, european eel, allis shad, zebrafish, striped catfish, mexican tetra, atlantic cod, medaka, northern pike, eastern mudminnow, brown trout, and rainbow trout), and 4 model species as evolutionary outgroups (human, mouse, fruitfly and roundworm) (Figure 1). Second, it supplies unbiased gene expression across 11 tissues using existing RNA-seq datasets (15) and up-to-date genomes. Indeed, due to the lack of reference genomes, previous RNA-seq data (15) were mapped on *de novo* contigs making expression quantification and cross-species investigations strongly biased. Finally, by combining genomic and remapped transcriptomic data, FEVER offers the possibility to rapidly explore gene expression evolution following speciation and duplication events. FEVER is implemented in R and is accessible at https://fever.sk8.inrae.fr/ with all major browsers. This website is free and open to all users and there is no login requirement.

**Figure 1:**
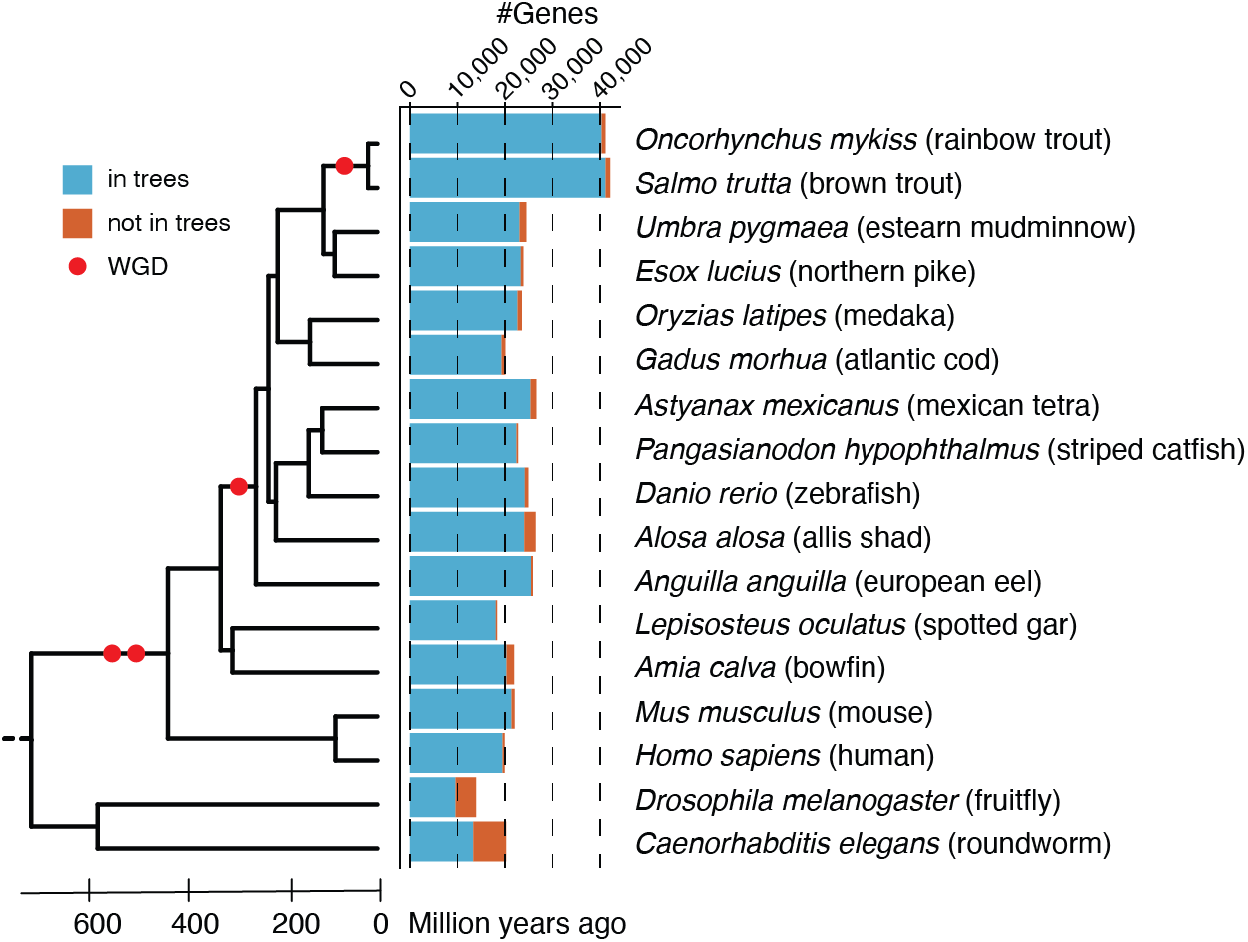
Species used for the reconstruction of gene trees. The Phylogenetic tree and divergence times are based on TimeTree (19) (v5) (http://www.timetree.org). Red dots indicate Whole Genome Duplication events (WGD). The bar plot shows the number of protein-coding genes of each species considered for the reconstruction of gene trees. Blue and red bars correspond to genes in trees and not in trees, respectively.

## MATERIALS AND METHODS

### Reconstruction of gene trees

To investigate the evolutionary history of extant genes across fishes, we reconstruct reconciled trees providing orthologous and paralogous relationships across 13 species covering the major fish lineages (rainbow trout, brown trout, eastern mudminnow, northern pike, medaka, atlantic cod, mexican tetra, striped catfish, zebrafish, allis shad, european eel, spotted gar, and bowfin) and 4 model species as evolutionary outgroups (human, mouse, fruitfly and roundworm) (Figure 1; Supplementary Table S1). First, the longest transcripts of all genes from the 17 aforementioned species are selected from GFF annotation files using the gstf_preparation.py program of the GeneSeqToFamily workflow (GeneSeqToFamily (16), parameters: --gff3 –fasta --headers -o --of --ff -l) and translated into proteins using the EMBOSS Transeq program (v6.6.0, https://emboss.sourceforge.net/, parameters: default). Next, all proteins are aligned to each other using an ALL vs ALL diamond search (diamond (17), v2.0.8, parameters: -k 0 -- outfmt 6 --evalue 1e-5) and clustered together based on alignment scores (hcluster_sg, http://treesoft.svn.sourceforge.net/viewvc/treesoft/branches/lh3/hcluster,v0.5.1-2, parameters: -m 750 -w 0 -s 0.34) to build families composed of homologous genes (orthologous and paralogous). Multiple alignments are then performed between all proteins composing each family (T-coffee (18), v11.00, parameters: -type=PROTEIN -method mafftgins_msa, muscle_msa,kalign_msa). Next, TreeBeSt is used to reconstruct codon-based multiple nucleotide alignments based on previous multiple protein alignments and coding sequences (TreeBeSt, https://github.com/Ensembl/treebest, v1.9.2, parameters: backtrans). Based on those alignments and the known species tree (TimeTree (19), http://www.timetree.org, v5), TreeBeSt (TreeBeSt, https://github.com/Ensembl/treebest, v1.9.2, parameters: best) reconstructs gene trees that are bootstrapped (100 times), reconciled with the species tree and rooted by minimizing the number of duplications and losses. Finally, those trees provide orthologous and paralogous genes across 17 species as well as their evolutionary history (speciation and duplication events). This methodology is strongly inspired by landmark comparative resources (12–14). Using this methodology, we generated 15,919 gene trees that contain 94.1% of all extant genes from the 17 species studied (96.4% for fish species) (Figure 1; Supplementary Table S1; Supplementary Figure 1). We also observe that 85% of all extant genes are covered by the 7,829 biggest gene families, and that the median number of species covered per gene family in those families equals 16 (Supplementary Figure 1). Altogether, those measures illustrate a great coverage of genes and species in the trees.

### Gene expression quantification

RNA-seq datasets across 11 tissues (brain, gills, heart, muscle, liver, kidney, bones, intestine, embryo, ovary, and testis) and 13 species (rainbow trout, brown trout, eastern mudminnow, northern pike, medaka, atlantic cod, mexican tetra, striped catfish, zebrafish, allis shad, european eel, spotted gar, and bowfin) were downloaded from a previous study (15). Further details on those samples are available in the biosample and bioproject files deposited in SRA under the PhyloFish umbrella project. Raw reads with known 3’ adaptor and low-quality bases (Phred score < 20) were trimmed with TrimGalore (https://github.com/FelixKrueger/TrimGalore, v.0.6.6, parameters: -r_clip 13 - three_prime_clip 2). Next, we mapped the trimmed reads from each library against reference genomes and annotated transcripts using Salmon (20) (v.1.8, parameters: defaults). Gene-expression levels were measured in transcripts per kilobase million (TPM), a unit which corrects for both feature length and sequencing depth. Tissue-specificity indexes are based on the Tau metric of tissue specificity (21) ranging from 0 (broad expression) to 1 (restricted expression). Tau is calculated for each gene across brain, gills, heart, muscle, liver, kidney, bones, intestine, embryo, ovary and testis. Tau is not calculated when at least one expression value is missing. A gene is considered tissue-specific when its Tau is greater than 0.9. Tau is defined as:

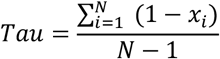

where N is the number of tissues and *x*_*i*_ is the expression profile component normalized by the maximal component value.

### FEVER web server implementation

FEVER is implemented in R (shiny) and is compatible with most web browsers (Google Chrome, Apple Safari, Mozilla Firefox, and Microsoft Edge) across the major operating systems (Windows, MacOS, and Linux). FEVER is hosted by the sk8 project which is based on technologies such as GitLab and CI/CD, Docker and Kubernetes. Using this website requires session cookies.

## RESULTS

FEVER provides a simple interface to investigate the evolution of gene expression across fish species. To enter a query gene, FEVER supplies a tool to search for genes of interest across 17 species (Figure 1) based on their gene ID, gene symbol or their longest transcript ID (Figure 2A). Based on this query gene, FEVER displays the reconstructed gene tree which provides orthologous and paralogous relationships across species through speciation and duplication events over evolution (Figure 2B). In addition, FEVER provides the expression (as z-scores on a heatmap) of all genes belonging to the tree across species and tissues (Figure 2C). The tissue specificity (Tau) of all genes, and the tissue a gene is specific to (when applicable) are displayed on the heatmap (Figure 2C). For an easier comparison across species and tissues, bar plots showing expression values (TPM) are also shown (Figure 2D). Gene trees and expression values can be downloaded from the web service. Gene trees, nucleotide and protein alignments have been deposited to Zenodo (https://doi.org/10.5281/zenodo.10844755).

**Figure 2:**
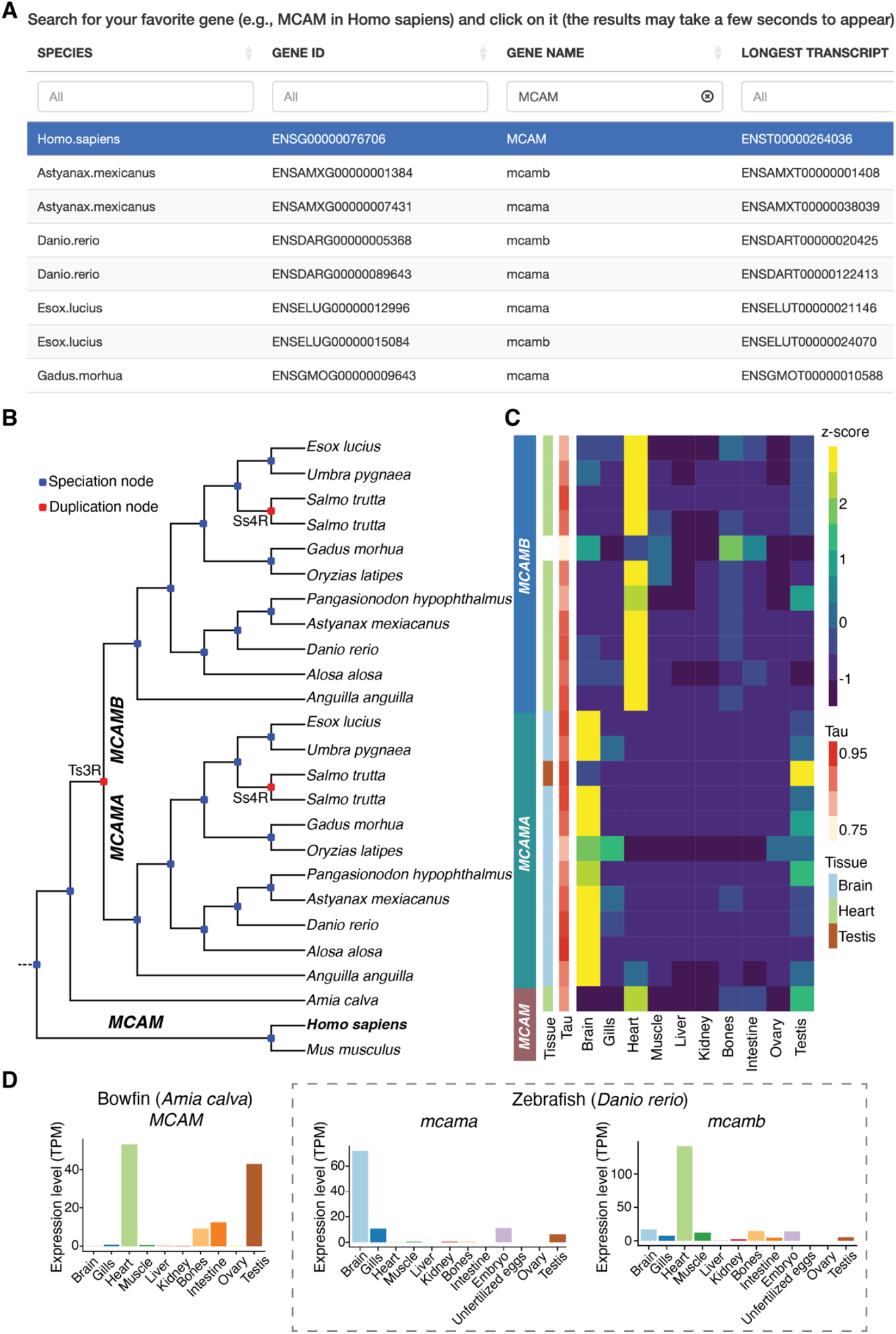
The FEVER web server interface and result page based on the query gene *MCAM* (Melanoma cell adhesion molecule). (A) FEVER provides an easy-to-use interface to search for genes of interest across 17 species. (B) FEVER displays the gene tree containing the query gene with duplication and speciation nodes annotated, in red and blue, respectively. (C) The heatmap shows gene expression across species and tissues as well as tissue-specificity indexes (Tau) and the tissue the genes are specific to, when applicable. (D) The bar plots show expression values (TPM) across tissues and organs for genes belonging to the same tree of the query gene. Expression values for *MCAM* in bowfin and *mcama*/*mcamb* in zebrafish are shown as a case study.

### Case study

To illustrate the features of FEVER, we provide and interpret the results of FEVER using *MCAM* as a query gene that is involved in cell adhesion and in cohesion of the endothelial monolayer at intercellular junctions in vascular tissue. FEVER revealed that this gene was duplicated by the teleost-specific WGD (Ts3R) and maintained in two copies (*MCAMA* and *MCAMB*) in most teleost species (Figure 2B). At the expression level, we note that *MCAMA* is specific to brain and *MCAMB* to heart in most teleost fish species, while the pre-Ts3R *MCAM* gene was likely to be particularly expressed in heart (Figure 2C), as inferred from *MCAM* expression in bowfin and mammals (from external data: https://apps.kaessmannlab.org/evodevoapp/). Additionally, *MCAMA* and *MCAMB* were duplicated once again by the salmonid-specific WGD (Ss4R) resulting in four *MCAM* copies in the brown trout, for instance. Interestingly, one post-Ss4R *MCAMA* paralog became specific to testis while the other one kept the pre-Ss4R *MCAMA* brain specificity, and the two post-Ss4R *MCAMB* paralogs maintained the pre-Ss4R *MCAMB* heart specificity. This example illustrates that gene tissue specificity may change following WGD events and that FEVER is a useful tool to assess gene evolutionary dynamics and identify potential sub/neo-functionalization. Expression values (TPM) can be easily compared across tissues as exemplified (Figure 2D) in bowfin (one *MCAM* copy) and zebrafish (two *MCAM* copies, *mcama* and *mcamb*).

## DISCUSSION

The FEVER web server supplies a simple interface to rapidly investigate the evolution of genes of interest across fish species following gene duplication events. It is noteworthy that one should be careful when inferring gene loss over evolution, because apparent gene loss can also be the result of missing genes in the annotations (e.g., mitochondrial genes) or inconsistent gene trees (e.g., fast-evolving genes). In this framework, it was previously observed that a fraction of gene trees reconstructed with similar methods in teleost fishes was inconsistent with their syntenic context resulting in misplaced WGD events (22). Consequently, SCORPiOs (22) was developed to correct those inconsistencies using synteny-based approaches. Thus, we plan to integrate SCORPiOs to our gene tree reconstruction pipeline in the next update. Furthermore, caution should be exercised when making cross-species comparisons of expression data in embryos and gonads, as synchronizing developmental stages and maturation status across species poses particular challenges. Future updates should also enable the user to blast any sequence of interest to investigate the evolution of the closest genes present in our dataset. Finally, we plan to progressively add additional species and expression data from other sources for teleost and non-teleost species.

## DATA AVAILABILITY

FEVER, gene trees, and computed expression values are freely accessible at https://fever.sk8.inrae.fr/ by all users without a login requirement. Gene trees, nucleotide and protein alignments have been deposited to Zenodo (https://doi.org/10.5281/zenodo.10844755).

## ACKNOWLEDGMENTS

We thank Yann Guiguen, Julien Bobe, Amaury Herpin, and Alexandra Depincé for discussions. FEVER is hosted by the sk8 project coordinated by INRAE.

---

**Supplementary Figure 1:**
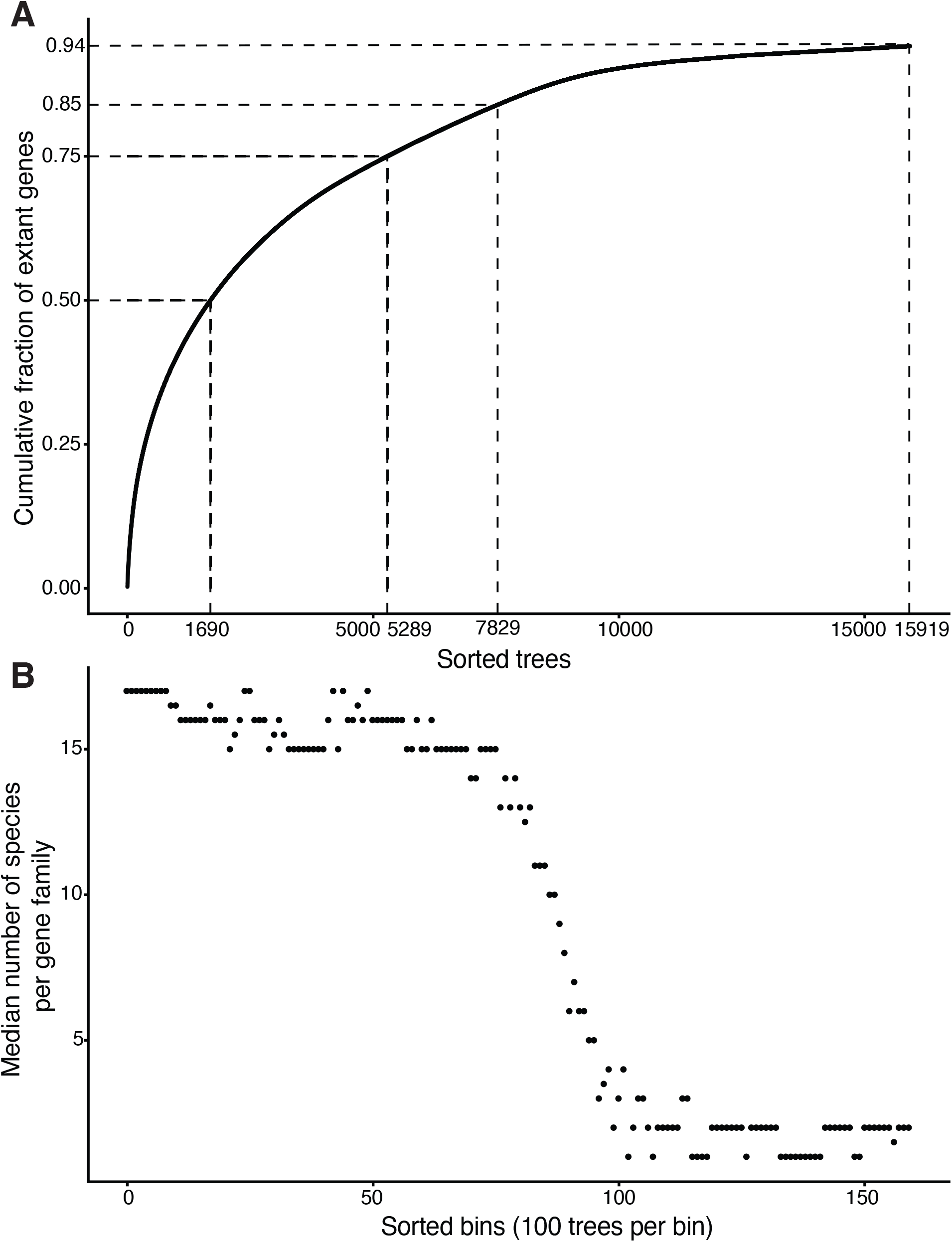
Characterization of gene families. (A) Cumulative fraction of all extant genes covered by gene families. Gene families are sorted decreasingly based on its number of genes on the x axis. (B) Median number of species per gene family. Gene families are sorted and binned (100 trees per bin) decreasingly based on its number of genes on the x axis.

**Supplementary Table S1:** Species and genomes used in this study.

| species scientific name | species common name | assembly | ncbi | Ensembl | #genes | #genes in trees | Fraction of genes in trees |
| --- | --- | --- | --- | --- | --- | --- | --- |
| Oncorhynchus mykiss | rainbow trout | USDA_OmykA_1.1 | GCF_013265735.2 | NA | 41123 | 40326 | 0.980619118 |
| Salmo trutta | brown trout | fSalTru1.1 | GCF_901001165.1 | NA | 42094 | 41149 | 0.977550245 |
| Umbra pygmaea | eastern mudminnow | ASM1680114v1 | GCA_016801145.1 | NA | 24516 | 23102 | 0.942323381 |
| Esox lucius | northern pike | Eluc_V3 | GCA_000721915.3 | ensembl V95 | 23875 | 23326 | 0.977005236 |
| Oryzias latipes | medaka | ASM223467v1 | GCA_002234675.1 | ensembl V95 | 23560 | 22658 | 0.961714771 |
| Gadus morhua | atlantic cod | gadMor1 | NA | ensembl V95 | 20093 | 19290 | 0.960035833 |
| Astyanax mexicanus | mexican tetra | Astyanax_mexicanus-2.0 | GCA_000372685.2 | ensembl V95 | 26584 | 25373 | 0.954446283 |
| Pangasianodon hypophthalmus | striped catfish | GENO_Phyp_1.0 | GCF_009078355.1 | NA | 22742 | 22441 | 0.986764577 |
| Danio rerio | zebrafish | GRCz11 | GCA_000002035.4 | ensembl V95 | 24909 | 24200 | 0.971536392 |
| Alosa alosa | allis shad | AALO_Geno_1.1 | GCA_017589495_2 | NA | 26440 | 24082 | 0.910816944 |
| Anguilla anguilla | european eel | fAngAng1.pri | GCF_013347855.1 | NA | 25888 | 25492 | 0.984703337 |
| Lepisosteus oculatus | spotted gar | LepOcu1 | GCA_000242695.1 | ensembl V95 | 18335 | 18059 | 0.984946823 |
| Amia calva | bowfin | AmiCal1 | GCA_017591415.1 | NA | 21948 | 20360 | 0.927647166 |
| Mus musculus | mouse | GRCm38.p6 | GCA_000001635.8 | ensembl V95 | 22040 | 21377 | 0.96991833 |
| Homo sapiens | human | GRCh38.p12 | GCA_000001405.27 | ensembl V95 | 19938 | 19495 | 0.977781121 |
| Drosophila melanogaster | fruitfly | BDGP6 | GCA_000001215.4 | ensembl V95 | 13903 | 9594 | 0.690066892 |
| Caenorhabditis elegans | roundworm | WBcel235 | GCA_000002985.3 | ensembl V95 | 20222 | 13323 | 0.65883691 |

## REFERENCES

1. Ohno, S. (1970) Evolution by Gene Duplication. Springer, New York.

2. Kaessmann, H. (2010) Origins, evolution, and phenotypic impact of new genes. Genome Res., 20, 1313–26.

3. Chen, S., Krinsky, B.H. and Long, M. (2013) New genes as drivers of phenotypic evolution. Nat. Rev. Genet., 14, 645–60.

4. Necsulea, A. and Kaessmann, H. (2014) Evolutionary dynamics of coding and non-coding transcriptomes. Nat. Rev. Genet., 15, 734–48.

5. Long, H.K., Prescott, S.L. and Wysocka, J. (2016) Ever-Changing Landscapes: Transcriptional Enhancers in Development and Evolution. Cell, 167, 1170–1187.

6. Brawand, D., Soumillon, M., Necsulea, A., Julien, P., Csárdi, G., Harrigan, P., Weier, M., Liechti, A., Aximu-Petri, A., Kircher, M., et al. (2011) The evolution of gene expression levels in mammalian organs. Nature, 478, 343–8.

7. Jaillon, O., Aury, J.-M., Brunet, F., Petit, J.-L., Stange-Thomann, N., Mauceli, E., Bouneau, L., Fischer, C., Ozouf-Costaz, C., Bernot, A., et al. (2004) Genome duplication in the teleost fish Tetraodon nigroviridis reveals the early vertebrate proto-karyotype. Nature, 431, 946–57.

8. Parey, E., Louis, A., Montfort, J., Guiguen, Y., Roest Crollius, H. and Berthelot, C. (2022) An atlas of fish genome evolution reveals delayed rediploidization following the teleost whole-genome duplication. Genome Res., 32, 1685–1697.

9. Berthelot, C., Brunet, F., Chalopin, D., Juanchich, A., Bernard, M., Noël, B., Bento, P., Da Silva, C., Labadie, K., Alberti, A., et al. (2014) The rainbow trout genome provides novel insights into evolution after whole-genome duplication in vertebrates. Nat. Commun., 5, 3657.

10. Xu, P., Xu, J., Liu, G., Chen, L., Zhou, Z., Peng, W., Jiang, Y., Zhao, Z., Jia, Z., Sun, Y., et al. (2019) The allotetraploid origin and asymmetrical genome evolution of the common carp Cyprinus carpio. Nat. Commun., 10, 1–11.

11. Parey, E., Louis, A., Montfort, J., Bouchez, O., Roques, C., Iampietro, C., Lluch, J., Castinel, A., Donnadieu, C., Desvignes, T., et al. (2023) Genome structures resolve the early diversification of teleost fishes. Science, 379, 572–575.

12. Muffato, M., Louis, A., Poisnel, C.-E. and Roest Crollius, H. (2010) Genomicus: a database and a browser to study gene synteny in modern and ancestral genomes. Bioinformatics, 26, 1119–21.

13. Nguyen, N.T.T., Vincens, P., Dufayard, J.F., Roest Crollius, H. and Louis, A. (2022) Genomicus in 2022: comparative tools for thousands of genomes and reconstructed ancestors. Nucleic Acids Res., 50, D1025–D1031.

14. Harrison, P.W., Amode, M.R., Austine-Orimoloye, O., Azov, A.G., Barba, M., Barnes, I., Becker, A., Bennett, R., Berry, A., Bhai, J., et al. (2023) Ensembl 2024. Nucleic Acids Res., 52, D891–D899

15. Pasquier, J., Cabau, C., Nguyen, T., Jouanno, E., Severac, D., Braasch, I., Journot, L., Pontarotti, P., Klopp, C., Postlethwait, J.H., et al. (2016) Gene evolution and gene expression after whole genome duplication in fish: The PhyloFish database. BMC Genomics, 17, 1–10.

16. Thanki, A.S., Soranzo, N., Haerty, W. and Davey, R.P. (2018) GeneSeqToFamily: a Galaxy workflow to find gene families based on the Ensembl Compara GeneTrees pipeline. Gigascience, 7, 1–10.

17. Buchfink, B., Xie, C. and Huson, D.H. (2015) Fast and sensitive protein alignment using DIAMOND. Nat. Methods, 12, 59–60.

18. Notredame, C., Higgins, D.G. and Heringa, J. (2000) T-Coffee: A novel method for fast and accurate multiple sequence alignment. J. Mol. Biol., 302, 205–17.

19. Kumar, S., Suleski, M., Craig, J.M., Kasprowicz, A.E., Sanderford, M., Li, M., Stecher, G. and Hedges, S.B. (2022) TimeTree 5: An Expanded Resource for Species Divergence Times. Mol. Biol. Evol., 39.

20. Patro, R., Duggal, G., Love, M.I., Irizarry, R.A. and Kingsford, C. (2017) Salmon provides fast and bias-aware quantification of transcript expression. Nat. Methods, 14, 417–419.

21. Yanai, I., Benjamin, H., Shmoish, M., Chalifa-Caspi, V., Shklar, M., Ophir, R., Bar-Even, A., Horn-Saban, S., Safran, M., Domany, E., et al. (2005) Genome-wide midrange transcription profiles reveal expression level relationships in human tissue specification. Bioinformatics, 21, 650–659.

22. Parey, E., Louis, A., Cabau, C., Guiguen, Y., Roest Crollius, H. and Berthelot, C. (2020) Synteny-Guided Resolution of Gene Trees Clarifies the Functional Impact of Whole-Genome Duplications. Mol. Biol. Evol., 37, 3324–3337.

